## Supplemental information for "Experimental nerve block study on painful withdrawal reflex responses in humans"

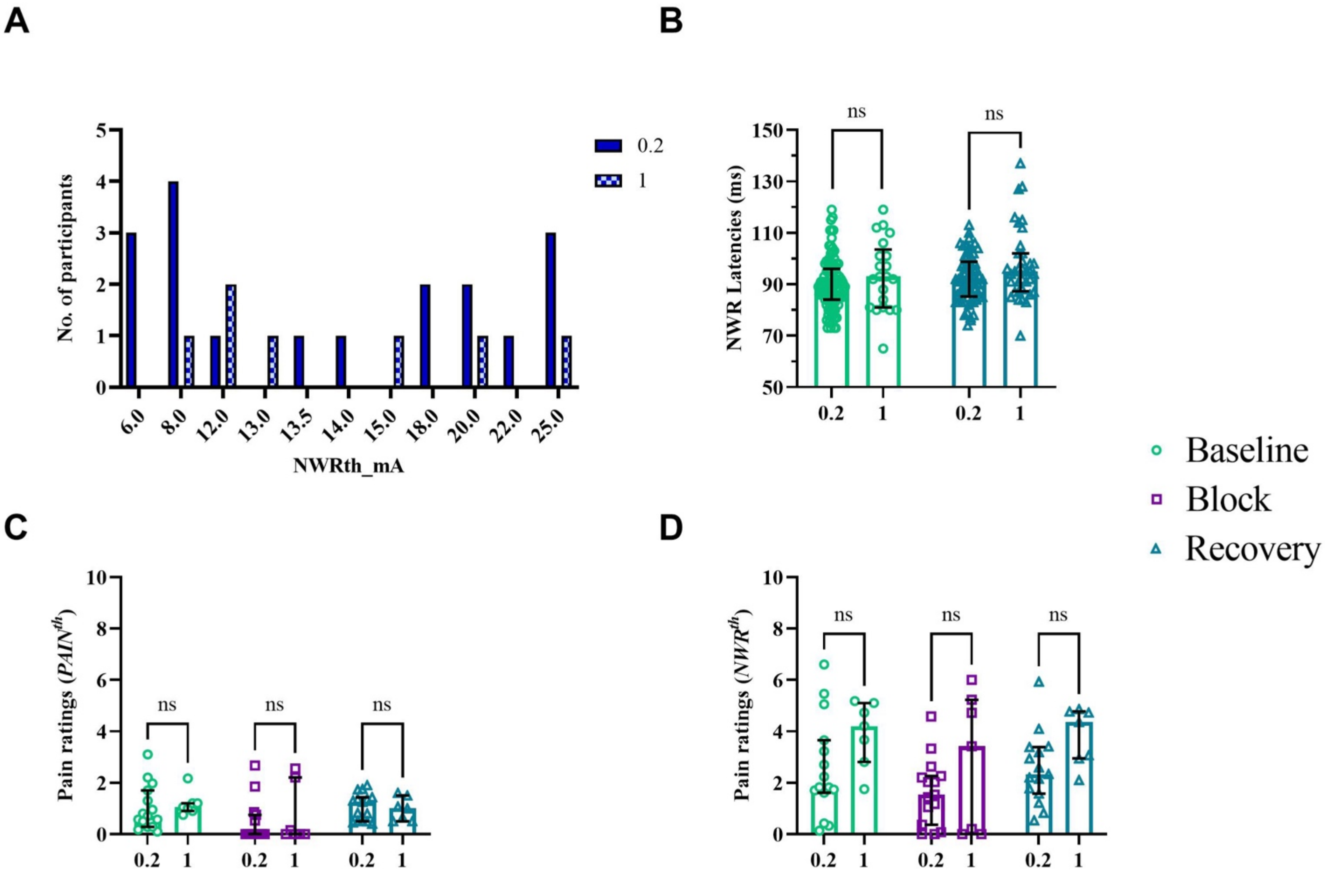


**Supplementary Figure 1. A. Distribution of *NWR$^{th}$* separated by pulse duration.** Those 7 subjects in whom a longer pulse duration (1.0 ms, checkered) was used had NWR thresholds at similar current intensities as the rest of the participants in whom a shorter pulse duration (0.2 ms, filled) was used. The y-axis shows the number of participants, and the x-axis shows the current intensities at which reflex thresholds were established. **B. NWR latencies separated by pulse duration.** There were no statistical differences in NWR latencies based on pulse duration (0.2 or 1.0 ms) within intact and recovery groups (**NWR latencies at 0.2 ms**: baseline 90.0 (12.0) ms, recovery 92.0 (16.5) ms; **NWR latencies at 1.0 ms:** baseline 93.0 (22.5) ms, recovery 95.0 (14.8) ms, f(3) = 9.72, p = 0.0211, post hoc test: p(baseline) = 0.384, p(recovery) = 0.194, n = 22). **C-D. Pain ratings at *PAIN$^{th}$* and *NWR$^{th}$* separated by pulse duration across conditions.** There were no statistical differences in pain ratings at *PAIN$^{th}$* or *NWR$^{th}$* based on pulse duration within baseline, block, and recovery groups (***PAIN$^{th}$* at 0.2 ms**: baseline 0.7 (1.4), block 0.0 (0.8), recovery 1.3 (0.9); ***PAIN$^{th}$* at 1.0 ms**: baseline 1.1 (0.3), block 0.0 (2.2), recovery 1.0 (1.0), f(5) = 2.38, p = 0.795, n = 22. Non-significant results in C are shown only for clarification. ***NWR$^{th}$* at 0.2 ms**: baseline 1.8 (2.0), block 1.5 (1.9), recovery: 2.3 (1.8); ***NWR$^{th}$* at 1.0 ms**: baseline 4.2 (2.3), block 3.4 (5.2), recovery

4.4 (1.8), f(5) = 13.77, p = 0.017, post hoc test: p(baseline) = 0.155, p(block) = 0.060, p(recovery) = 0.251, n = 22). Figs. B-D, Kruskal-Wallis test. ns = not significant.

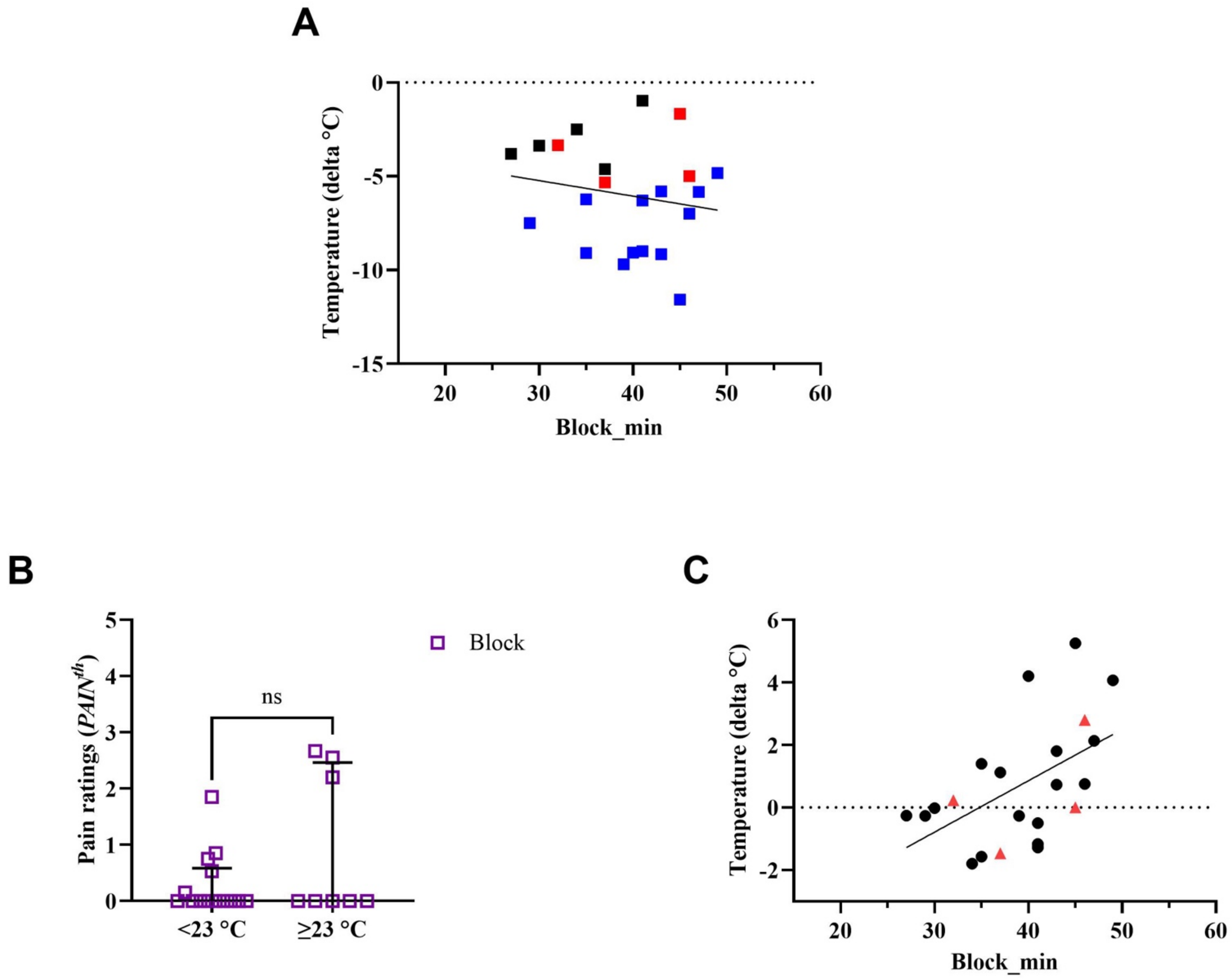


**Supplementary Figure 2. A. Comparison between the duration of nerve block and change in cooling sensitivity.** There was no association between nerve block duration and change in cold detection threshold (r = -0.167, p = 0.458, CI 95% [-0.559, 0.286], n = 22, Spearman r). Blue squares represent participants with a CDT below 23°C and black squares represent participants with a CDT above 23°C. Red triangles represent the 4 subjects with increased pain at *PAIN$^{th}$* during the block. **B. Pain ratings at *PAIN$^{th}$* between subjects with CDT below or above 23°C.** Pain ratings did not differ between subjects who had a CDT lower or higher than the lower end 95% CI of the reference data (Magerl et al., 2010) for CDT in the foot (pain ratings <23°C: 0.0 (0.6), pain ratings ≥23°C: 0.0 (2.5), p = 0.534, U = 47.50, n =

22, Mann-Whitney test; ns = not significant). **C. Comparison between the duration of nerve block and change in warming sensitivity.** A significant positive correlation was found between nerve block duration and warm detection threshold (r = 0.535, p = 0.010, CI 95 % [0.133, 0.786], n = 22, Spearman r). Red triangles represent the 4 subjects with increased pain at $PAIN^{th}$ during the block.

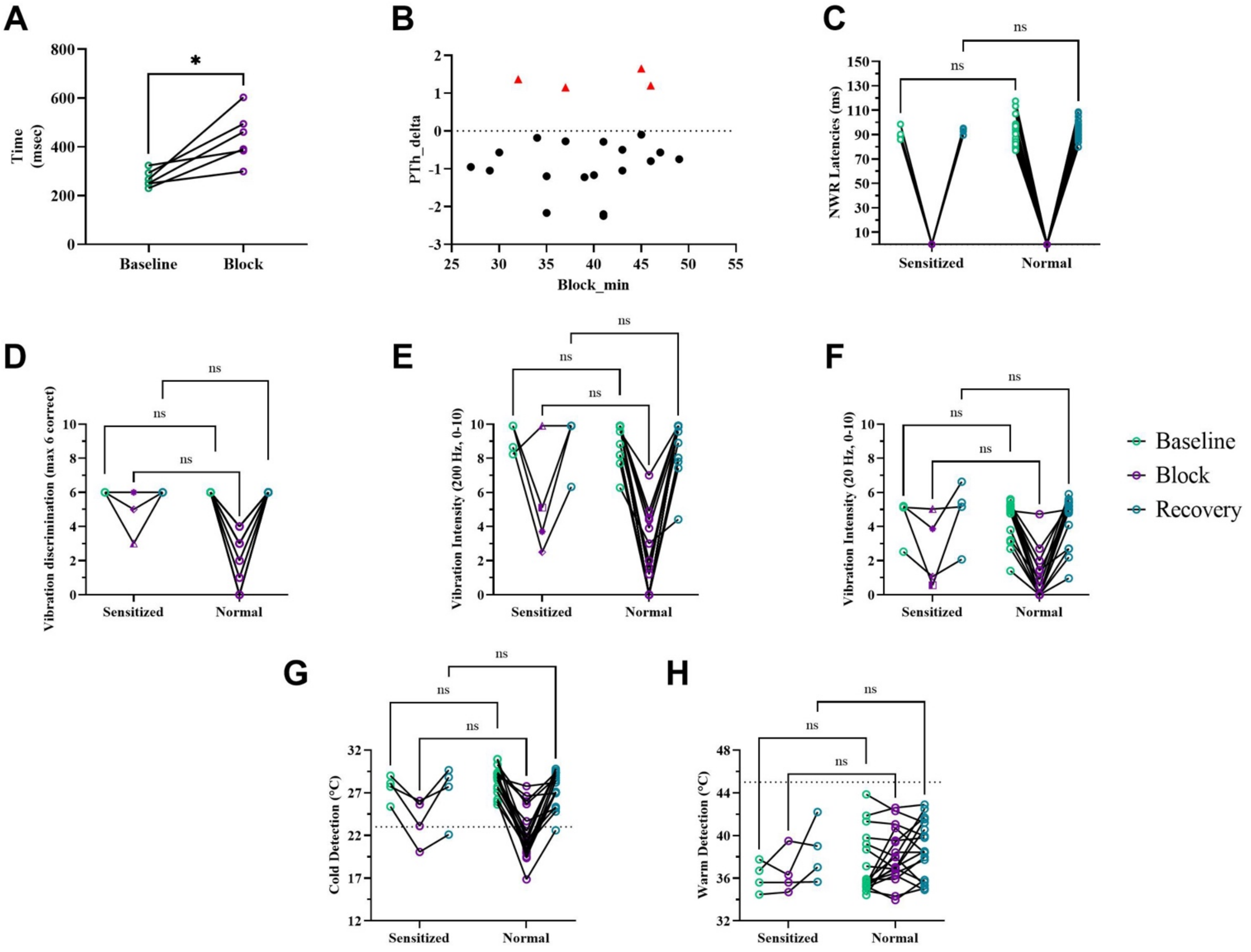


**Supplementary Figure 3. A. Reaction time increased during nerve block.** The median reaction time showed a significant increase during the block (baseline 258.8 (56.3) ms, block 426.2 (158.5) ms, W = 21, p = 0.031, n = 6, Wilcoxon test), suggesting a possible shift in conducting afferents. **B. Duration of nerve block.** There was no significant correlation between pain ratings at $PAIN^{th}$ and the nerve block duration (r = 0.151, p = 0.5025, CI [-0.301, 0.548], n = 22, Spearman r). Those 4 subjects who reported an increase in pain at $PAIN^{th}$ during the block are indicated by red triangles. **C. Comparison of reflex latencies between sensitized and normal subjects.** To simplify, the minority of subjects with increased pain at $PAIN^{th}$ during the block are categorized in the 'sensitized' group while the rest are categorized in the 'normal' group. NWR latencies did not differ between sensitized (n=4) and normal (n=18) subjects (**Sensitized**: baseline 89.6 (9.7) ms, block 0.0 (0.0) ms, recovery: 93.6 (4.8) ms; **Normal**: baseline 91.6 (12.0) ms, block 0.0 (0.0) ms, recovery 93.3 (12.8) ms, f(2) = 48.72, p < 0.001, post hoc test: p(baseline & recovery) > 0.999, n = 22). **D-F Comparison of performance on vibratory tests during the block between sensitized and normal groups.** Vibration intensity ratings in the blocked condition cannot explain why pain did not reduce in the sensitized group. Although one out of four subjects in the sensitized group did not show a reduction in vibration intensity ratings during the block (triangle-shaped, Figs. E & F), that same subject performed poorly on the three-alternative forced-choice detection task (3-AFC, Fig. D). The different shapes of data points (triangle, square, diamond, and star) in D-F represent the same 4 subjects for all vibration tests (**3-AFC Sensitized**: baseline 6.0 (0.0), block 5.0 (3.0), recovery 6.0 (0.0); **3-AFC Normal**: baseline 6.0 (0.0), block 2.5 (3.3), recovery 6.0 (0.0), f(5) = 57.63, p < 0.001, post hoc test: p(baseline & recovery) > 0.999, p(block) = 0.337, n = 21. Multiple participants have overlapping values in D). **200 Hz Sensitized**: baseline 9.3 (1.6), block 4.4 (5.9), recovery 9.9 (2.7); **200 Hz Normal**: baseline 9.9 (1.3), block 0.6 (4.0), recovery 9.9 (1.2), f(5) = 41.69, p < 0.001, post hoc test: p(baseline & recovery) > 0.999, p(block) = 0.418, n = 22. **20 Hz Sensitized**: baseline 5.1 (2.0), block 2.5 (4.1), recovery 5.3 (3.5); **20 Hz Normal**: baseline 5.0 (1.6), block 0.0 (1.4), recovery: 5.0 (1.7), f(5) = 36.16, p < 0.001, post hoc test: p(baseline & recovery) > 0.999, p(block) = 0.533, n = 22). **G-H. Comparison of performance on thermal tests during the block between sensitized and normal responses.** The 4 subjects in the sensitized group did not have different cold and warm detection thresholds (CDT and WDT) in the blocked condition from those in the normal group. The dotted lines represent lower (CDT) or higher (WDT) limits for normal threshold values in the lower limb (**CDT Sensitized**: baseline 27.9 (2.8)°C, block 24.4 (5.1)°C, recovery 28.3 (6.0)°C; **CDT Normal**: baseline 28.8 (1.9)°C, block 21.3 (4.1)°C, recovery 27.7 (2.3)°C, f(5) = 34.35, p < 0.001, post hoc test: p(baseline, block & recovery) > 0.999, n = 22. **WDT Sensitized**: baseline 36.2 (2.7)°C, block 36.0 (3.7)°C, recovery 38.0 (5.3)°C; **WDT Normal**: baseline 35.8 (4.1)°C, block 37.5 (3.4)°C, recovery 38.5 (5.5)°C, f(5) = 6.981, p = 0.221, n = 22. Non-significant results in H are shown only for clarification). Figs. C-H, Kruskal-Wallis test. ns = not significant.

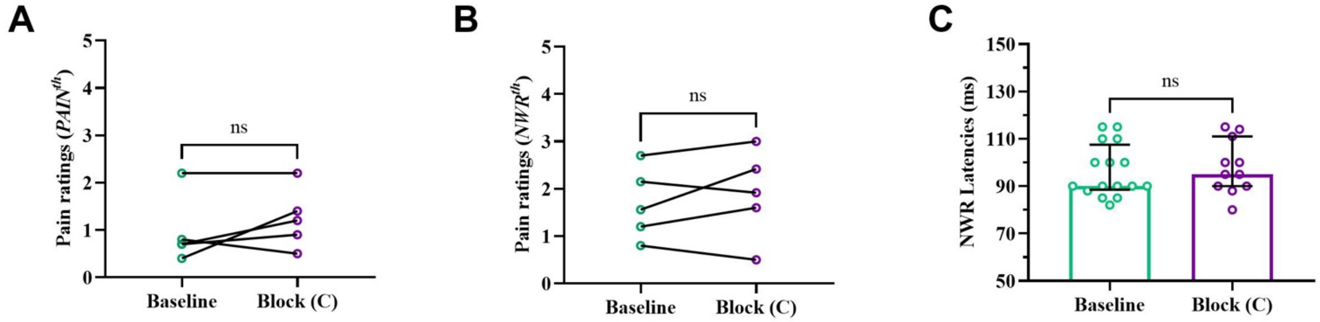


**Supplementary Figure 4. Pain and reflex measurements on the intact test site with a nerve block applied to the contralateral leg**. There were no statistical differences in pain ratings on the intact test site during the contralateral block (***PAIN^th***: baseline 0.7 (1.0), block 1.2 (1.1), W = 6, p = 0.375, n = 5; ***NWR^th***: baseline 1.6 (1.4), block 1.9 (1.7), W = 8, p = 0.375, n = 5, Wilcoxon test; Figs. A-B). Reflex latencies on the intact site were unaffected during the contralateral block (baseline 90.0 (19.0) ms, block 95.0 (21.0), W = 77.5, p = 0.615, n = 27, Mann-Whitney test; Fig. C). C = contralateral.
